# Dual neutralization of influenza virus hemagglutinin and neuraminidase by a bispecific antibody leads to improved antiviral activity

**DOI:** 10.1101/2023.03.16.532941

**Authors:** Romila Moirangthem, Sapir Cordela, Dina Khateeb, Michal Mandelboim, Friederike Jönsson, Timothée Bruel, Yotam Bar-On

## Abstract

Targeting multiple viral proteins is pivotal for sustained viral suppression. In recent years, several broadly neutralizing antibodies that target the influenza virus hemagglutinin and neuraminidase glycoproteins have been developed. However, the impact of dual neutralization of these two glycoproteins on the course of infection has not been thoroughly tested. Here we demonstrate that a bispecific antibody that neutralizes both the hemagglutinin and the neuraminidase has a dual antiviral activity as it blocks infection and prevents the release of progeny viruses from the infected cells. We further show that dual neutralization of the hemagglutinin and the neuraminidase by a bispecific antibody is advantageous over antibody combination as it results in an improved neutralization capacity and augmented antibody effector functions. Notably, the bispecific antibody showed enhanced antiviral activity in influenza virus-infected mice. These findings suggest that dual neutralization of the hemagglutinin and neuraminidase could be effective in controlling influenza virus infection.

## Introduction

Most vaccines in clinical use today prevent infection because of their ability to elicit neutralizing antibodies that block pathogen entry^1^. However, highly mutable viruses such as influenza virus and HIV-1 pose a great challenge for vaccine development, as attempts to design a vaccine that will elicit neutralizing antibodies against all the circulating variants have yet to be successful^2^. As an alternative, broadly neutralizing antibodies (bNAbs) that are able to recognize and neutralize a large portion of the circulating viruses were developed, and passive immunization with bNAbs was shown to suppress influenza virus and HIV-1 replication in preclinical and human studies^3–8^. Nevertheless, since highly mutable viruses have shown the capacity to develop resistance to bNAbs during the course of antibody therapy^6–8^, passive immunization with a combination of broadly neutralizing antibodies directed against different viral epitopes is currently being tested for its ability to better control these viruses^9^ and was also a leading therapeutic strategy in the COVID-19 pandemic^10^. Moreover, combination therapy has become a cornerstone of cancer treatment^11^.

Most of the bNAbs against influenza virus target the influenza virus surface glycoprotein hemagglutinin (HA)^12,13^, which is responsible for the binding of the virus to cells and for initiating the infection^14^. The HA can be divided into two domains: the head, which comprises the receptor-binding site that binds sialic acids on the surface of the target cells, and the stem, which is responsible for fusion of the viral membrane and the cellular membrane^15^. bNAbs against the HA can both target the HA head and inhibit the attachment of the virus to the target cell or the HA stem and interfere with the viral fusion machinery^12,13,16^. Despite the unique breath of these broadly neutralizing anti-HA antibodies, several viral mutations have been reported to impair the antibody binding and function. For example, the HA mutation A388V was shown to impair the binding of bNAbs that bind the HA stem, and head mutations that can make the virus completely resistant to strain-specific antibodies can also enable the virus to escape bNAbs that target the same region^17,18^. The other major surface glycoprotein of influenza virus, the neuraminidase (NA), has been recently suggested as a promising target for bNAbs^19^. NA is an enzyme that cleaves the sialoside receptor from the cell surface and enables the progeny virus to be released from the infected cell during viral budding^20^. As the NA exhibits a slower antigenic drift than the HA, broadly neutralizing antibodies against this glycoprotein usually show broader cross-reactivity^19^.

Of note, the effect of dual antibody targeting, in which both the HA and the NA are neutralized, has not been thoroughly tested. This is of special interest since it has been shown that a functional balance between HA and NA is required for efficient viral replication^21–23^. For example, reverse genetics studies showed that reduced fitness of influenza virus that resulted from an increased HA affinity could be restored by an increased NA activity ^21–23^. Moreover, it was shown that mutations in the HA protein can lead to secondary mutations in the NA protein, and it has been suggested that balance between the HA and NA activity has a great impact on viral growth and viral fitness^21–23^. Finally, it was recently shown that neutralization by anti-HA stem antibodies can also block the NA activity^24^. This was attributed to steric inhibition of the viral neuraminidase (NA) by the Fc region of anti-HA stem antibodies and provided another example for the close proximity of the viral NA and HA^24^.

The unique interplay of the HA and NA glycoproteins, and our previous findings in which we demonstrated the use of dual antibody therapy for sustained viral suppression of highly mutable viruses^9^ prompt us to perform in-depth analysis of dual antibody-mediated neutralization of the influenza virus HA and NA. Of note, a combination of broadly neutralizing antibodies was recently shown to be effective in both protecting and treating various viral infections including hepatitis B virus^25^, HIV-1^3–6^ and SARS-CoV-2^10^ in both preclinical trials and in the clinic. The fact that antibody combination therapy was shown to be safe, tolerable and was able to limit the emergence of the antibody-resistant viruses, has drawn much interest in using such approach for better controlling influenza virus infections, in which current therapeutics show limited success^13^. Moreover, the emergence of flu pandemic that occur every few decades stresses the importance of developing novel therapeutic approaches for severe and deadly flu infections^26^. Here we generated a bispecific antibody that neutralize the HA and NA of various influenza virus strains and we have demonstrated the bispecific antibody has a dual antiviral activity and an improved neutralization activity. Moreover, we evaluate the therapeutic capacity of the bispecific antibody in an animal model and the beneficial effects of neutralizing the HA and NA with a bispecific antibody in comparison with a dual antibody administration.

## Results

To provide in-depth analysis of the virological and immunological outcomes of dual neutralization of the influenza virus hemagglutinin (HA) and neuraminidase (NA), we have designed and generated a bispecific antibody (BsAb) that neutralizes both of these viral glycoproteins. For the generation of the bispecific antibody, we have used combination of the stem-directed anti-HA broadly neutralizing antibody CR6261^27^ and the recently isolated anti-NA broadly neutralizing antibody 1G01^19^. These antibodies were selected based on their exceptional breadth and potency^19,28^. Plasmids encoding the light chains and heavy chains of these antibodies with engineered C_H_3 domains that promotes heterodimerization^29^ were used to transfect Expi-CHO cells and the bispecific was purified from the cell supernatant (Figure 1A). Following successful purification of the BsAb we infected MDCK cells with influenza virus A/Puerto Rico/8/1934 (H1N1) and stained uninfected cells and influenza virus infected cells with the BsAb 48 hrs following the infection. A significant increase in the BsAb staining was seen in the infected MDCK cells in comparison with uninfected cells (Figure 1B). Next, we tested if the BsAb can bind to influenza virus virions. For this we have coated EL4 cells with influenza virus A/Puerto Rico/8/1934 (H1N1). EL4 cells cannot be infected with this virus, but instead the virus binds the cell membrane of the EL4 cells^30^. Thus, following incubation with the virus, the cells are coated with the viral particles. Influenza virus-coated cells were positively detected by the BsAb (Figure 1B). Thus, the BsAb could recognize the HA and NA glycoproteins on the membrane of the influenza virus-infected cells and on H1N1 virions. To further evaluate the binding of the BsAb to the HA and NA, we have directly tested the binding of the BsAb to these glycoproteins in ELISA. ELISA plates were coated with the HA glycoprotein from different subtypes of Influenza A virus, H1N1 (A/California/04/2009), Influenza A H1N1 (A/Puerto Rico/8/1934), Influenza A H5N1 (A/Vietnam/1194/2004) and were stained with BsAb or anti-HA CR6261 antibody. Similarly, NA glycoproteins of H1N1 (A/California/04/2009) and Influenza A H5N1 (A/Hubei/1/2010) were stained with the BsAb or anti-NA antibody 1G01. In agreement with previous works ^13,19^, the monoclonal antibodies CR6261 and 1G01 successfully recognized the tested HA and NA glycoproteins (Figure 1C-D). Of note, the BsAb showed significant binding to the HA and NA glycoproteins and the binding was comparable to the binding of CR6261 and 1G01(Figure 1C-D). Furthermore, the binding of the BsAb significantly increased in wells in which when both the HA and the NA proteins were present, indicating that both arms of the bispecific are engaged with the two antigens (Figure 1E).

**Figure 1.**
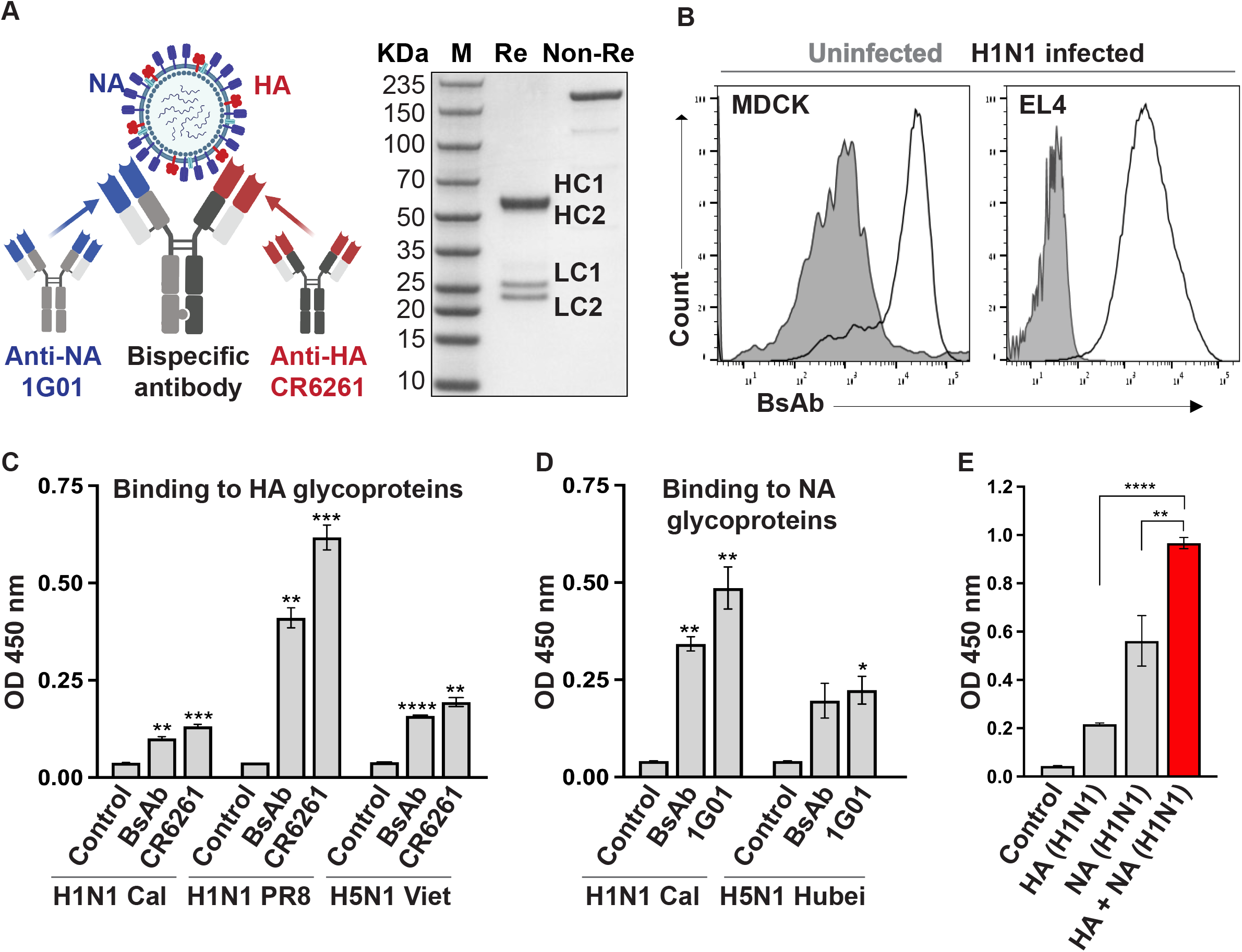
Binding of the bispecific antibody to the HA and NA glycoproteins. (A) Schematic representation of the BsAb and SDS-PAGE analysis. The antibody was isolated from the cell culture supernatant by protein G antibody purification and the purified bispecific antibody was analyzed on SDS-PAGE. Re=reducing conditions, Non-Re=non-reducing, HC= Heavy, LC=light chain. (B) FACS staining of infected cells and influenza virus-coated cells. The left histogram shows the binding of the bispecific antibody to virions-coated cells, and the right histogram shows the binding to influenza virus-infected cells (H1N1 A/PR/8/34). Grey histograms depict the binding of the bispecific antibody to uncoated or uninfected cells. The figure shows representative staining. Four independent experiments were performed. Chain conditions (C-E) ELISA plates were coated with 0.5 μg of HA glycoproteins, 0.5 μg of NA glycoprotein or 0.5 μg of a combination of HA+NA (0.25 μg each) and stained with the bispecific antibody (BsAb), CR6261 or 1G01. HA and NA were derived from different influenza virus strains (indicated in the x-axis). The y-axis depicts the O.D. levels. Data is derived from triplicates. Representative results from three independent experiments are shown. Statistically significant differences are shown (student’s t test, *p<0.05, **p<0.01, ***p<0.001, ****p<0.0001). Shown are mean values and error bars.

We have design the BsAb to have a dual antiviral activity, which we termed ‘block and lock’. This was based on the notion that blockade of the HA by BsAb will neutralize the influenza virus HA and will prevent the infection of the target cells, while neutralization of the NA will prevent the release of progeny viruses from the infected cells. To test the neutralization capacity of the BsAb we have incubated influenza virus A/Puerto Rico/8/1934 (H1N1) with serial dilutions of BsAb, CR6261 or 1G01 and evaluated the ability of the antibodies to neutralize the infection of MDCK cells in a microneutralization assay^31^. While both the CR6261 and the 1G01 antibodies were able to neutralize influenza virus and block the infection of MDCK cells, the BsAb demonstrated a significantly higher neutralization capacity with an average half maximal inhibitory concentration (IC_50_) of 0.3 μg/ml in comparison the CR6261 IC_50_=1.51 μg/ml and 1G01 IC_50_=1.9 (Figure 2A). We postulated that the dual neutralization of the HA and NA result in augmented neutralization due to the involvement of the NA in facilitating the infection of the target cells in addition to its role in the release of progeny viruses (^32^, Figure 2A). This is in line with the viral neutralization that we observed when 1G01 antibody was used to block infection (Figure 2A). Next, we have compared the neutralization capacity of the BsAb and a combination of CR6261 and 1G01 antibodies. Strikingly, the neutralization capacity of the BsAb (IC_50_=0.51 μg/ml) was superior to a combination of CR6261 and 1G01 (IC_50_=0.92 μg/ml, Figure 2B). To determine whether treatment of influenza virus-infected cells with the BsAb will reduce the release of progeny viruses from the infected cells. Infected-MDCK cells were treated with the BsAb 24 hrs after infection was established and viruses from the cell supernatant were collected from the antibody-treated cells and untreated cells 48 hrs following infection. The collected viruses were then used to infect fresh MDCK cells and the level of infection was evaluated after 48 hrs. Viruses that were collected from infected untreated MDCK cells resulted in a robust infection of the freshly thawed MDCK cells while minimal or no infection was seen when viruses were collected from BsAb-treated infected cells (figure 2C). We concluded that the BsAb shows high neutralization capacity and is also capable of preventing the budding of newly formed viruses from infected cells.

**Figure 2.**
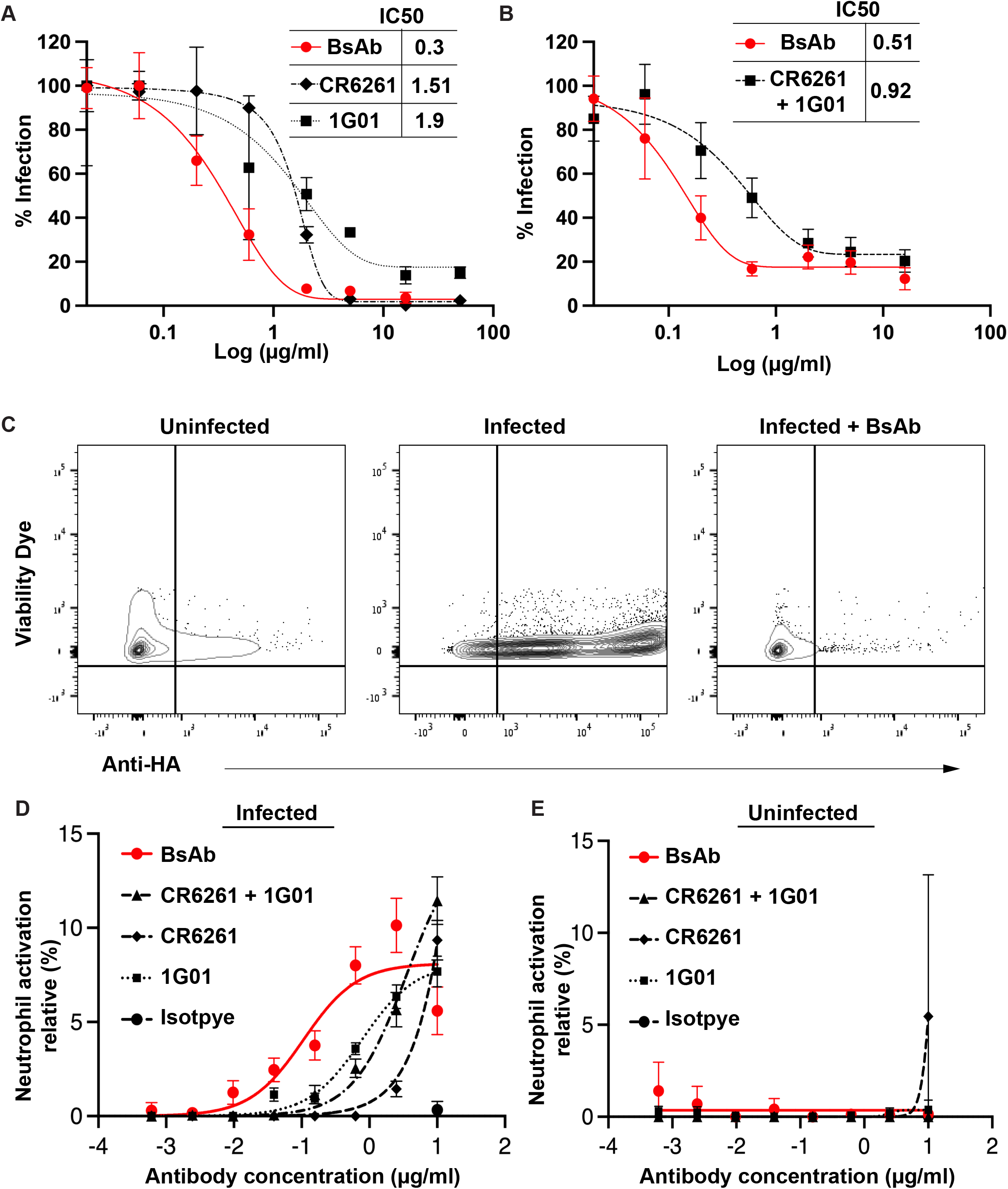
The dual antiviral activity of the BsAb. (A-B) A/Puerto Rico/8/1934 influenza virus was incubated with the CR6261 antibody, with 1G01 antibody, with the BsAb antibody or a combination of CR6261 and 1G01. MDCK infection was evaluated by analyzing the expression of the influenza virus nucleoprotein in the infected cells. To calculate the % infection, the results were compared to cells that were infected with influenza virus in the absence of antibodies. Antibody concentrations are indicated in the x-axis. A summary of three independent experiment is shown. (C) Plots that depict the percentage of infected cells (HA positive cells) in MDCK cells that were incubated with supernatant from infected cells (middle panel) or incubated with supernatant from infected cells that were treated with the bispecific antibody (right panel). The number in each panel depicts the percentage of influenza virus-infected cells. One representative experiment is shown out of three performed. (D-E) % of neutrophil activation following incubation of infected (left graph) and uninfected (right graph)MDCK cells with BsAb, CR6261, 1G01, or a combination of CR6261+1G01. The antibody concentration is depicted in the x axis. Neutrophile activation was measured by analyzing CD62L shedding and CD11b upregulation after subtraction of the values that were observed with no target cells but with the addition of the antibodies.

We postulated that improved neutralization of the bispecific antibody compared to a combination of the CR6261 and 1G01 antibodies could be due to steric inhibition caused by anti-stem HA antibodies that block access to the influenza NA glycoprotein as shown by Kosik et al.^24^and others^33^. Since such steric inhibition is mediated mainly by the fragment crystallizable region (Fc region) of the anti-HA antibody ^24^ we next aimed to determine if any differences in the antibody Fc-mediated effector functions will be observed when the HA and NA are targeted by the BsAb or by a combination of CR6261 and 1G01. We focused our analysis on the ability of the antibodies to engage the human Fc receptor CD32a since it was previously shown to play a major role during influenza virus infection^34^. Mouse neutrophiles that were engineered to express the human CD32a were co-cultured with A/Puerto Rico/8/1934 (H1N1)-infected MDCK cells that were incubated with various amounts of the BsAb, CR6261, 1G01 or a combination of CR6261 and 1G01 and the levels of neutrophile activation was measured by analyzing CD62L shedding and CD11b upregulation (Figure 2D-E). Incubation of the infected cells with the BsAb has led to significantly higher percentage of neutrophile activation in comparison with CR6261, 1G01 or a combination of CR6261 and 1G01 across different antibody concentrations (Figure 2D). Moreover, no neutrophile active activation was observed toward uninfected MDCK cells, indicated that the neutrophile activation is mediated by the binding of the antibody to HA and NA of the surface of the infected cells (Figure 2E).

Based on the dual function of the BsAb, its enhanced neutralization capacity and its augmented Fc-mediated functions (Figure 2), we next evaluated the therapeutic capacity of the BsAb. For through *in vivo* comparison of the additive effect of dual neutralization of the HA and NA over single antibody neutralization, we first follow the course of influenza infection following administration of a single neutralizing antibody. Administration of CR6261 at 15 mg/kg was previously shown to prevent mice weight loss and to significantly improve mice survival of influenza virus-infected mice^35^. Thus, we first followed the course of the infection following administration of lower doses of CR6261. At a dose of 6 mg/kg CR6261 was still able to effectively prevent the mice weight loss and 85% of the mice survived the infection (Supplementary Figure 1). However, at 2 mg/kg CR6261 failed to prevent the mice weight loss and only 30% of the mice survived the infection (Supplementary Figure 1). Thus, we choose the 2 mg/kg antibody dosage for comparing the antiviral activity of CR6261, 1G01 and the BsAb.

C57BL/6 mice were administrated with 2mg/kg of CR6261, 1G01 and the BsAb. Four hours following the antibody administration the mice were infected with 4.6 x 10^3^ PFU of A/Puerto Rico/8/1934 (H1N1). In accordance with our *in vitro* findings (Figure 1-2), administration of the BsAb significantly prevented the mice weight loss in comparison with mice that were administrated with either CR6261 or 1G01 (Figure 3A). Moreover, the viral loads of A/Puerto Rico/8/1934 (H1N1) were measured in the mice lungs three days following infection of untreated mice or of mice that were administrated with CR6261, 1G01 and the BsAb. We observed a significant reduction in the lung viral loads in all of the antibody-treated mice, however the viral loads in the BsAb treated mice (VL= 48564 copies/ml) were significantly lower than in the CR6261 (VL=57329 copies/ml) or 1G01 (VL=49789 copies/ml) treated mice (Figure 3B). Since we demonstrated that administration of the BsAb can lead to reduced weight loss and viral loads compared to CR6261 and 1G01 (Figure 3A-B), we performed additional experiments in which we evaluated the course of the disease following administration of BsAb, CR6261,1G01 or a combination of CR6261 and 1G01. Only mice that were administered with the BsAb showed a significant reduction in the maximal weight loss following infection in comparison with untreated infected mice (Figure 3C). Importantly, the BsAb significantly improved the mice survival in comparison with CR6261 or 1G01 (Figure 4D). The BsAb also improved the mice survival compared to a combination of CR6261 and 1G01, albeit to a lesser extent (Figure 4D). We concluded that dual neutralization of the HA and NA by a bispecific antibody can improve antibody-based control of influenza virus infection.

**Figure 3.**
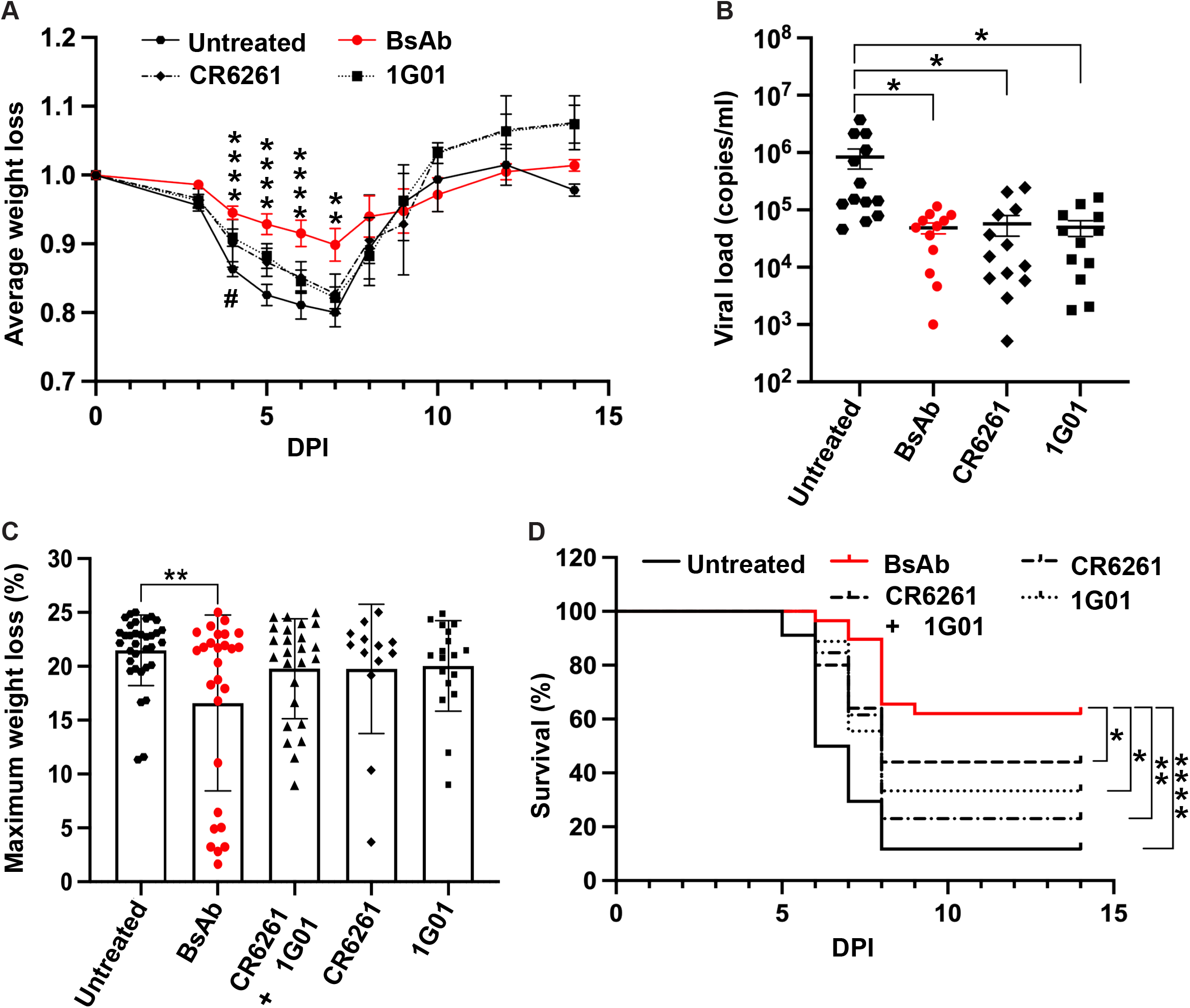
The antiviral activity of the BsAb in comparison with CR6261, 1G01 and antibody combination. (A-B) Mice weight loss (A) and viral titers in the lungs (B) were evaluated following infection with the A/Puerto Rico/8/1934 (4.6 x 10^3^ PFU) and administration of 2 mg/kg of the BsAb, CR6261, 1G01 or in untreated mice. Antibodies were administered 4 hours before infection by intraperitoneal injection. Statistically significant differences are indicated (*p < 0.05, **p < 0.01, **** p<0.0001). In (B), each data point depicts an individual mouse. Mean values and SD are shown in (A-B). In (A) the x axis indicates DPI (days post infection). The data is a summary of three independent experiments. (C-D) Maximal mice weight loss (C) and % survival following infection with the A/Puerto Rico/8/1934 (4.6 x 10^3^ PFU) and administration of 2 mg/kg of the BsAb, CR6261, 1G01, a combination of CR6261 and 1G01 or in untreated mice. Antibodies were administered 4 hours before infection by intraperitoneal injection. In (C) maximal weight loss was calculated by dividing the minimal weight of each mouse with the weight of each mouse at day 0. Each data point depicts an individual mouse. Shown are mean values and error bars. Statistically significant differences are indicated (**p < 0.01). In the survival curve (D), the number of mice in each group is n=22 and mice losing 25% of their initial weight were considered as dead. The statistical differences between BsAb and other antibodies treated conditions were determined by Gehan-Breslow-Wilcoxon test (*p<0.05, **p<0.01, ****p<0.0001).

## Discussion

In recent years, combination therapy has revolutionized cancer therapy and is one of the most promising approaches that is currently being tested in clinical trials for the treatment of highly diverse viruses^10,11^. Passive immunization with a combination of broadly neutralizing antibodies directed against different viral epitopes has already showed to effectively suppress highly mutable viruses such as HIV-1^9^ and this approach is also of great interest since it is a leading therapeutic strategy for COVID-19^10^. Unlike other enveloped RNA viruses, such as HIV-1 and SARS-CoV-2 that contain only one dominant surface protein, influenza virus virions and influenza virus-infected cells express high levels of two distinct glycoproteins, the HA and the NA^36^. The accessibility of these viral glycoproteins on virions and on influenza virus-infected cells place them as possible target for antibody combination therapy. There is currently high need for novel therapeutic for controlling influenza infection as the efficacy of the flu vaccine is suboptimal and drug resistant to the commonly used drug oseltamivir carboxylate (Tamiflu) is frequent^37–39^. Moreover, thus far the use of mRNA-based vaccines that showed high efficacy against SARS-CoV-2 had not significantly improved the protection from influenza virus infection^1^.

In this work we uncovered several advantages of dual antibody neutralization of the influenza virus HA and NA. First, as the HA viral glycoprotein is responsible for viral entry to the target cells and the NA facilitate the budding of the newly formed viruses from the infected cells, we show that the BsAb antibody can inhibit the viral entry to the target cells and the viral release from the target cells. Second, we demonstrated that dual neutralization of the HA and NA results in an improved viral neutralization compared to neutralization of only the HA glycoprotein, which is considered the dominant viral protein that facilitate viral infection. We postulate that the enhanced neutralization activity of the BsAb is due to multifactional role of the NA in influenza virus life cycle^20,23,32^. As the role of the NA in releasing to progeny viruses from the infected epithelial cells is well documented and described, additional studies have suggested that NA may play other roles in the viral attachment and entry to the target cells^24,32^. In agreement with this, we and others demonstrated that neutralizing antibodies against the NA can significantly impair viral infection^31^. Given the unique interplay of the HA and NA^23^, it would be intriguing to identify combinations of HA and NA antibodies with synergetic activity, in which escape mutations that arise in one glycoprotein can render the other protein more susceptible to antibody neutralization or impairs its activity.

One clear advantage of using a bispecific of using a BsAb for neutralizing the HA and NA over a combination of two monoclonal antibodies is that the generation of a BsAb is more economical^40^. Nevertheless, this work highlights functional differences in the activity of BsAb that target the HA and NA compared to antibody combination. In particular, we observed that the BsAb has an improved neutralization activity and an augmented ability to engage Fc receptor in comparison with a combination of the CR6261 and 1G01 antibodies. The reasons for these differences are not clear. Since the CR6261 antibody is directed against the stem of the HA, and because the Fc portion of anti-stem antibodies was shown to sterically inhibit the NA^24^, we hypothesis that a possible explanation for the improved activity of the BsAb over CR6261 and 1G01 is steric inhibition of the NA by CR6261, which hinders the binding of 1G01 to the NA. Thus, other combination of anti-HA and anti-NA antibodies could show better antiviral activity. Alternatively, these differences could also be attributed to a more synchronized neutralization of the HA and NA by the BsAb in comparison with a combination of CR6261 and 1G01, however we did not address this possibility and it should be further evaluated in additional experiments.

Viral inhibition by drug combination has shown to be effective against highly mutable viruses and has dramatically changed the outcome for people living with HIV-1^3,6,10,41,42^. In addition, Antibody combination therapy has been extensively studied against HIV-1, HBV and SARS-CoV-2^9^. However, a through evolution of dual antibody neutralization of the HA and NA during influenza virus infection has not been performed despite the initial finding by Marathe et al. that a combination of anti-HA and anti-NA antibodies improve the survival of immunosuppressed mice following lethal infection with influenza B virus^43^. The recent isolation of neutralizing neuraminidase antibodies with exceptional breadth to multiple different influenza A and B virus from an H3N2 infected donor, has paved the way for effective neutralization of the NA and for a combination therapy in which both the HA and the NA are neutralized^19^. Out of the different NA antibodies that were isolated, the 1G01 antibody has the highest number of somatic hypermutations exhibit the broadest binding^19^, which prompt us to combine this antibody with CR6261 that binds the hemagglutinin of most group 1 influenza viruses. Dual inhibition of the HA and NA can also be achieved by antiviral drugs^37^. As anti-NA inhibitors are commonly used in the clinic for treating influenza virus infection^37^, our results suggest that the combination of such drugs with antibodies that neutralized the influenza virus HA could lead to improve the control of seasonal influenza virus infections and possible future influenza virus pandemics. Further studies are required to identify the HA and NA epitopes that can be targeted by such combined therapy in order to reduce the emergence of resistant viral strains.

## Materials and methods

### Cells, viruses and bacterial strains

The cell lines used in this study were the human alveolar basal epithelial cells (A549), the Madin-Darby canine kidney (MDCK) cells, T lymphoblast EL4 cells and primary mouse neutrophiles that were engineered to express the human CD32a. Cells were cultured in Dulbecco’s Modified Eagle Medium (DMEM) supplemented with 4500 mg/l D-glucose, 4 mM L-glutamine, 110 mg/l sodium pyruvate, 10% FBS, 1% penicillin–streptomycin, and 1% nonessential amino acids (NEAA). Cells were maintained at 37°C with 5% CO2. DH5α bacteria (Thermo Fisher Scientific,18258012) grown in LB media at 37°C were used for cloning and amplification of plasmid DNA for mammalian cell transfection. ExpiCHO™ Expression System Kit (Cat. No. A29133) was used for generating the monoclonal antibodies. The influenza viruses A/Puerto Rico/8/34 (H1N1) was used for the *in vitro* and *in vivo* experiments.

### Flow cytometry

For influenza virus infection, cells were incubated for 1 hr with A/Puerto Rico/8/34 (H1N1) at 37°C in a 6-well plate. Cells were cultured in serum-free media together with trypsin type IX (Sigma, T-0303). The cells were then washed and incubated with Dulbecco’s Modified Eagle Medium (DMEM) supplemented with 4500 mg/l D-glucose, 4 mM L-glutamine, 110 mg/l sodium pyruvate, 10% FBS, 1% penicillin–streptomycin, and 1% nonessential amino acids (NEAA). Staining was performed 48 hrs following infection. For coating of EL4 cells with A/Puerto Rico/8/34 (H1N1), cells were incubated with the virus for 4 hrs at 37°C. For staining of the infected cells were stained with 1 μg of CR6261, 1G01 or the BsAb for 1 hr at 4°C. Cells were then washed and incubated with the appropriate secondary antibody for 45 min. staining was evaluated using the BD LSRFortessa flow cytometer. Further analysis of the staining results was done using FlowJo software v10.7. Viability dye (65-0865-14) was used in order to gate on live cells only.

### ELISA

To test the binding of CR6261, 1G01 and the BsAb to the HA and NA glycoproteins, ELISA was performed by coating high-binding 96-well plates (Thermo scientific, Ca t#44-2404-21) with he glycoproteins (1xPBS with 0.05% Tween-20) and incubated with 200 μl/well blocking buffer (1xPBS with 2% skim milk) for 2 hours at room temperature. ELISA plates were coated with 0.5 μg of HA glycoproteins, 0.5 μg of NA glycoprotein or 0.5 μg of a combination of HA+NA (0.25 μg each) Immediately after blocking, plates were incubated with the appropriate antibody (CR6261, 1G01 or BsAb) for 1 hr. After washing 6 times with washing buffer, plates were incubated with anti-mouse IgG secondary antibody conjugated to horseradish peroxidase (HRP) diluted in blocking buffer at 1:10,000 dilution. Plates were developed by the addition of the HRP substrate, TMB (Life technologies, Cat #002023). The absorbance was immediately measured at 450 nm with an ELISA microplate reader (Infinite M200 PRO). Data were analyzed using Prism V8.0.

### Microneutralization assay

The neutralizing activity of CR6261, 1G01 the BsAb or a combination of CR6261, 1G01 were evaluated using previously described protocol^31^. To calculate the % infection, the results were compared to cells that were infected with influenza virus in the absence of antibodies. MDCK cells were used as target cells. The virus was preincubated with the antibody and the mixture was added to a monolayer of MDCK cells (70–80% confluent in 96-well plates). The concentrations of the antibodies are indicated in the figure. After incubation at 37 °C for 1 hr, the cell monolayer was washed three times with PBSX1 and re-incubated for 18–20 h at 37 °C with medium (DMEM supplemented with 50 U ml^-1^ penicillin, 50 μg ml^-1^ streptomycin, 25 mM HEPES and 1 μg/ml TPCK-treated trypsin) containing monoclonal antibodies (at equivalent concentrations as during the virus co-incubation). Cells were fixed with 80% (v/v) acetone, blocked with 5% (w/v) non-fat milk diluted in PBS for 30 min at room temperature, and quenched with 3% (v/v) hydrogen peroxide (in PBS) by incubating for a further 20 min at room temperature. Cells were stained with biotinylated anti-NP antibody (EMD Millipore; 1:2,000), followed by HRP-conjugated streptavidin (Jackson Immunoresearch; 1:5,000). Plates were developed by the addition of the HRP substrate, TMB (Life technologies, Cat #002023)The absorbance was immediately measured at 450 nm with an ELISA microplate reader (Infinite M200 PRO). Data were analyzed using Prism V8.0.

### CD32a activation

Primary neutrophils from mice humanized for CD32a were incubated with MDCK cells 48 hrs after infection with A/Puerto Rico/8/34 (H1N1) and addition of CR6261, 1G01, a combination of CR6261+1G01 or the BSAb. The amount of the antibodies is indicated in the x-axis. After 45 min of culture the levels of CD62L and CD11b were evaluated by flow cytometry in order to analyze the levels of neutrophil activation. The levels of CD62L and CD11b seen in neutrophils incubated with the antibodies, but in the absence of MDCK cells, was used to calculate the background staining levels. Uninfected MDCK cells that were incubated with similar amounts of antibodies were used as an additional control.

### NA inhibition assay

To test the ability of the BsAb to impair the release of newly formed viruses from the infected cells by blocking the NA, A549 cells were infected with A/Puerto Rico/8/1934 (H1N1), and 48 hours post infection, the bispecific antibody was added to the cell culture. Three days after the infection, the cell supernatant was collected and was used to infect freshly thawed A549 cells after removal of remaining antibodies from the media. Supernatant from infected cells that were not treated with the antibody was used as control. Infection was evaluated 48 hrs later by staining to the influenza virus HA using the CR6261 antibody and calculating the percentage of the live HA^+^ cells. Viability dye (65-0865-14) was used in order to gate on live cells only. Staining was evaluated using the BD LSRFortessa flow cytometer. Further analysis of the staining results was done using FlowJo software v10.7.

### Mice infection and antibody administration

Female C57BL/6 mice (8 weeks old) mice were administrated with 2 mg/kg or 6 mg/kg of CR6261, 1G01, a combination of CR6261+1G01 and the BsAb (intraperitoneal injection). Antibodies were diluted in PBSX1 and were analyzed on SDS-PAGE before injection. Four hours following the antibody administration the mice were infected with 4.6 x 10^3^ PFU of A/Puerto Rico/8/1934 intranasally. The virus weas diluted in PBSX1 in a volume of 25 μl. Antibody levels were verified by bleeding the mice 1 day post antibody administration and by testing the antibody levels in ELISA. The mice weight was monitored daily and mice that lost more than 20% of their initial weight were scarified. For viral load measurement, the mice lungs were harvested, and the amount of the virus was evaluated by real-time PCR as previously described^44^ and a standard curve was generated with known copies of viral RNA to calculate the copies/ml. maximal weight loss was calculated by dividing the minimal weight of each mouse with the weight of each mouse at day 0.

### Statistical Methods

Student’s t test and Gehan-Breslow-Wilcoxon test (for the survival experiments) were used to determine significant differences.

## Acknowledgments

We thank J. Yewdell and Ivan Kosik for their help with designing the BsAb. This study was supported by the Israel Ministry of Science and Technology MOST: France-Israel Joint Projects (0005201), The U.S.-Israel Binational Science Foundation (BSF, 2021068) and the Michigan-Israel partnership.

