## Supplementary Figure 1 for "Dual neutralization of influenza virus hemagglutinin and neuraminidase by a bispecific antibody leads to improved antiviral activity"

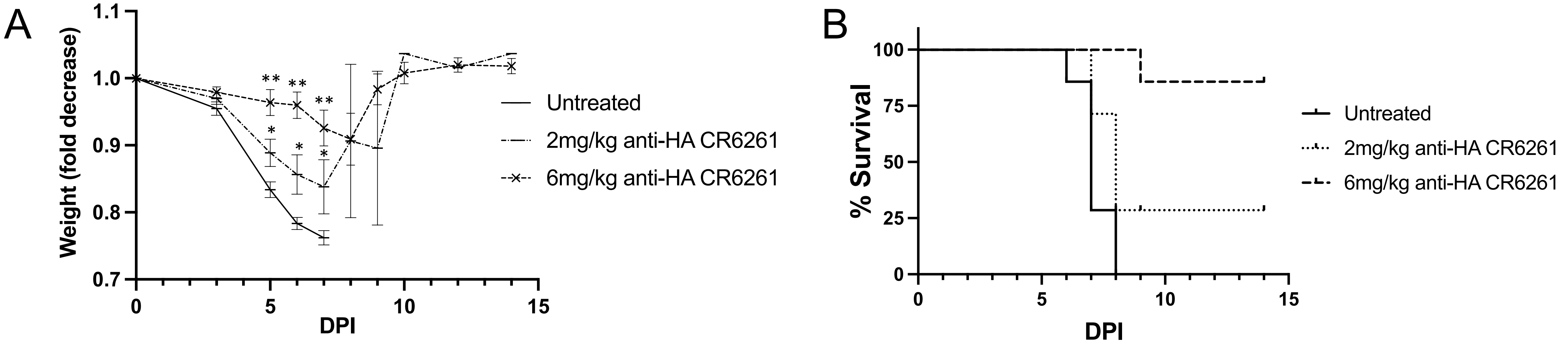


**Supplementary Figure 1. Mice infection and CR6261 administration. (**A-B) Mice weight loss (A) and survival (B) were evaluated following infection with the A/Puerto Rico/8/1934 (4.6 x 103 PFU) and administration of 2 mg/kg or 6 mg/kg of CR6261mice. Antibodies were administered 4 hours before infection by intraperitoneal injection. Statistically significant differences are indicated ($*p < 0.05$, $**p < 0.01$).
